# In Utero Alcohol Exposure Reprograms Mammary Gland Development and Stem/Progenitor Cell Activity in a Non-Tumorigenic Mouse Model

**DOI:** 10.64898/2026.09.22.753591

**Authors:** Zhikun Ma, Amanda B Parris, Evelyn I Espinoza, Xiaohe Yang

**Author notes:** **Correspondence:** Xiaohe Yang Ph.D.

## Abstract

In utero alcohol exposure (IUE) can alter mammary development and increase mammary tumor susceptibility in experimental models. However, most previous studies have relied on carcinogen-induced or genetically tumor-prone models, leaving it unresolved whether prenatal alcohol exposure produces mammary developmental and molecular changes before the introduction of an oncogenic challenge. Here, we addressed this question using non-tumorigenic C57BL/6 mice. Pregnant dams received a Lieber–DeCarli liquid diet containing 1.7% (v/v) ethanol or an isocaloric control diet from gestational day 11 to 19, and mammary glands from female offspring were examined during puberty (postnatal day 40, PND40) and young adulthood (PND70). At PND40, IUE enhanced mammary ductal morphogenesis, increased terminal end bud numbers, and elevated epithelial proliferation. These developmental alterations persisted at PND70, with increased lateral branching and proliferative activity. IUE also altered mammary epithelial composition, expanding the basal/myoepithelial compartment and the CD24^high^CD49f^high^ mammary repopulating unit/stem cell-enriched population. Mammary epithelial cells from IUE-exposed offspring exhibited increased colony-forming cells (CFCs), mammosphere formation, and 3D Matrigel growth, indicating enhanced stem/progenitor-associated activity. At the molecular level, IUE increased ERα expression and phosphorylation together with Cyclin D1, c-Myc, E2F1, and Bcl-2, and enhanced AKT, ERK, and STAT3 phosphorylation, IL-6 expression, and SOX2 at PND40. Importantly, major components of this signaling phenotype remained altered at PND70, including ERα-associated signaling, AKT/ERK activation, and IL-6/STAT3 signaling, whereas the pubertal increase in SOX2 was not sustained. Together, these findings demonstrate that prenatal alcohol exposure induces profound and persistent histomorphological, cellular, and signaling alterations in the mammary gland in the absence of a carcinogenic challenge or genetically introduced oncogenic driver. These results identify intrinsic developmental reprogramming of normal mammary tissue as a potential basis through which prenatal alcohol exposure may modify subsequent responses to tumor-promoting or other environmental challenges.

## 1. Introduction

Breast cancer is a multifactorial disease arising from interactions among genetic susceptibility, hormonal regulation, lifestyle factors, and environmental exposures [1, 2]. Increasing evidence suggests that breast cancer susceptibility may also be influenced by exposures occurring much earlier in life, including during prenatal development. This concept is consistent with the developmental origins of health and disease (DOHaD) paradigm, in which environmental perturbations during critical developmental windows can induce persistent changes in tissue organization and regulatory programs that influence disease susceptibility later in life [3]. The mammary gland may be particularly sensitive to such developmental programming because mammary epithelial development begins in utero and is followed by extensive hormonally regulated growth and remodeling during puberty, pregnancy, lactation, involution[4–6]. Consequently, developmental exposures that alter mammary epithelial organization, cellular hierarchy, or signaling responsiveness could have long-lasting consequences for mammary gland biology and subsequent disease susceptibility [7, 8].

Alcohol represents an important developmental exposure in this context. In addition to being an established risk factor for breast cancer when consumed during adulthood [9], ethanol is a potent developmental toxicant. Prenatal alcohol exposure can disrupt fetal development through multiple mechanisms, including oxidative stress, generation of reactive metabolites, altered cellular signaling, and epigenetic regulation, producing developmental abnormalities collectively recognized as fetal alcohol spectrum disorders (FASD)[10–14]. Although the neurological consequences of prenatal alcohol exposure have received the greatest attention, alcohol can also perturb the development and long-term function of non-neural tissues [15–17]. These observations raise the broader question of whether prenatal alcohol exposure can durably reprogram mammary gland development and thereby modify susceptibility to mammary disease later in life.

Experimental studies provide evidence supporting this possibility. Prenatal alcohol exposure has been shown to increase susceptibility to chemically induced mammary tumors in rodent models, with alterations in tumor multiplicity, latency, and tumor phenotypes [18, 19]. Importantly, these tumor-associated effects are preceded or accompanied by changes in mammary gland development, including increased terminal end buds (TEBs), altered ductal morphogenesis, and dysregulation of hormone- and growth factor-responsive pathways [18, 19]. Changes in circulating estradiol, estrogen receptor (ER) signaling, insulin-like growth factor signaling, and epigenetic regulation have also been implicated [18, 20, 21]. Collectively, these findings suggest that prenatal alcohol exposure may establish persistent cellular and molecular changes in mammary tissue that subsequently modify its response to carcinogenic or oncogenic stimuli.

Mammary stem and progenitor cells provide a potential cellular basis for such developmental reprogramming. Mammary epithelial development and remodeling depend on dynamically regulated stem/progenitor-enriched populations that generate and maintain basal/myoepithelial and luminal epithelial compartments [4, 22–24]. These populations are particularly important during periods of active morphogenesis, including pubertal ductal growth, when highly proliferative TEBs drive epithelial expansion [25]. Alterations in the abundance, self-renewal capacity, differentiation, or regulatory signaling of mammary stem/progenitor populations can therefore produce persistent changes in mammary epithelial architecture and function and may influence subsequent susceptibility to neoplastic transformation [22–24]. The reported effects of prenatal alcohol exposure on TEB development and epithelial composition raise the possibility that mammary stem/progenitor cells are important targets of IUE-induced developmental programming.

An important unresolved question, however, is whether prenatal alcohol exposure establishes persistent changes in mammary development and epithelial regulation that may alter how the gland subsequently responds to tumor-promoting or other environmental influences. Most previous studies have relied on carcinogen-induced or genetically tumor-prone models [18, 20, 26]. These studies provide important proof-of-concept evidence that prenatal alcohol exposure can modify mammary tumor risk, but the presence of an exogenous carcinogenic challenge or pre-existing oncogenic driver makes it difficult to determine whether developmental alterations are already present before a tumor-promoting challenge and how they may subsequently interact with oncogenic signals. Our previous work using the MMTV-erbB2 transgenic mouse model further illustrated this issue. In that model, low-dose IUE altered mammary morphogenesis and significantly increased mammary tumor multiplicity, accompanied by enhanced ERα signaling and strong activation of ErbB2/EGFR-associated pathways, including AKT and ERK signaling [27]. IUE also altered mammary epithelial cell populations and increased the MaSC-enriched mammary repopulating unit (MRU) population, supporting a connection between developmental exposure, stem/progenitor cell regulation, and later tumor susceptibility [27]. These findings raised the question of whether similar developmental, cellular, and signaling alterations are present in normal mammary tissue in the absence of an oncogenic driver such as ErbB2.

Addressing this question requires examination of prenatal alcohol exposure in a non-tumorigenic mammary gland, where developmental phenotypes can be separated from oncogene-driven or carcinogen-induced effects. In the present study, we therefore used C57BL/6 mice to determine whether IUE alters mammary development and mammary epithelial stem/progenitor-associated activity in the absence of an oncogenic driver. We examined mammary morphogenesis and epithelial proliferation during puberty and young adulthood, characterized mammary epithelial subpopulations and stem/progenitor-associated functions, and evaluated hormone-responsive and growth/survival signaling pathways implicated by our previous tumor-model studies. We show that IUE accelerates mammary ductal morphogenesis and epithelial proliferation, alters mammary epithelial hierarchy and enhances stem/progenitor-associated activity, and induces persistent activation of ERα-, AKT/ERK-, and IL-6/STAT3-associated signaling. These findings demonstrate that prenatal alcohol exposure can induce persistent developmental and molecular alterations in the mammary gland in the absence of a pre-existing oncogenic driver, providing a cellular and signaling context through which developmental alcohol exposure may influence subsequent mammary disease susceptibility.

## 2. Materials and Methods

### 2.1 Antibodies

Antibodies against ERα (sc-8002), ERβ (sc-390243), ERK2 (sc-1647), Cyclin D1 (sc-246), Bcl-2 (sc-7382), AKT1 (sc-5298), E2F1 (sc-251), and β-actin (sc-47778) were obtained from Santa Cruz Biotechnology (Santa Cruz, CA, USA). Antibodies against p-ERα (Ser118; 2511), PR (8757), c-Myc (5605), p-AKT (Ser473; 4060), p-ERK1/2 (Thr202/Tyr204; 9101), IL-6 (D5W4V), SOX2 (D1C7J), p-STAT3 (Tyr705; D3A7), and STAT3 (124H6) were obtained from Cell Signaling Technology (Beverly, MA, USA). For immunohistochemistry, the Ki67 (PA5-19462), Alpha-Smooth Muscle Actin (α-SMA) Antibody (14-9760-82) were purchased from Invitrogen. Flow cytometry antibodies against CD16/32 (553141), CD49f (555735), and CD24 (553260) were purchased from BD Biosciences, and CD31 (102508), CD45 (103106), and Ter-119 (116208) were purchased from BioLegend.

### 2.2 Animals and in utero alcohol exposure

C57BL/6 mice were obtained from Jackson Laboratory (Bar Harbor, ME, USA). The mice were maintained in a temperature-controlled facility with a 12 h light–dark cycle. Except during the ethanol exposure period, mice were fed estrogen-free AIN-93G semipurified diet (Bio-Serv., Flemington, NJ, USA) ad libitum. All experimental procedures were approved by the Institutional Animal Care and Use Committee.

Breeding pairs were established at 8 weeks of age. Pregnant dams were randomly assigned to either the control or ethanol-exposure group on gestational day (GD) 10. From GD 11 to GD 19, dams in the ethanol group received a Lieber–DeCarli ’82 liquid diet containing 1.7% (v/v) ethanol, prepared daily by mixing one part 5% (v/v) ethanol liquid diet with two parts isocaloric control diet. Control dams received the corresponding isocaloric liquid control diet during the same period.

Each experimental group included 10 pregnant dams. Only female offspring were used for subsequent analyses. To minimize potential litter effects, no more than one female offspring from each litter was included in a given analysis. A total of 20 mice per group were used for various analyses, including whole-mount analysis, histopathology, Western blotting and flow cytometry, as described in the respective assay protocols. Mammary gland development and associated cellular and molecular changes were evaluated at postnatal day (PND) 40, representing the peripubertal stage, and PND70, representing young adulthood.

### 2.3 Mammary gland whole-mount preparation and morphometric analysis

At the indicated endpoints, fourth and fifth mammary glands were harvested and fixed overnight in Carnoy’s fixative. Tissues were subsequently rehydrated, stained with carmine alum as previously described [28], dehydrated through graded alcohols, and cleared in xylene. Whole-mount preparations were digitally imaged for morphological analysis.

At PND40, pubertal mammary morphogenesis was evaluated by quantifying the number of terminal end buds (TEBs) per gland. Mammary glands from five mice per experimental group were analyzed. At PND70, ductal complexity was assessed by quantifying lateral branching per 10 mm² of mammary tissue.

### 2.4 Immunohistochemistry and immunofluorescence

Mammary gland tissues were fixed in 10% neutral-buffered formalin, processed using standard histological procedures, embedded in paraffin, and sectioned for staining.

For chromogenic immunohistochemistry (IHC), sections were deparaffinized and rehydrated, followed by heat-induced antigen retrieval in 0.01 M citrate buffer (pH 6.0). Endogenous peroxidase activity was quenched with 3% H₂O₂ for 10 min. Sections were incubated overnight at 4 °C with primary antibodies against phosphorylated estrogen receptor-α (p-ERα; 1:100) or Cyclin D1 (1:100). After washing, sections were incubated with the appropriate biotinylated secondary antibody, followed by signal amplification using the VECTASTAIN® Elite ABC kit (Vector Laboratories). Immunoreactivity was visualized using diaminobenzidine (DAB), and sections were counterstained with hematoxylin. Images were acquired using a Nikon light microscope under identical imaging conditions within each experiment.

For immunofluorescence analysis, tissue sections were blocked with 10% goat serum in PBST and incubated overnight at 4 °C with primary antibodies against Ki67 (1:1000) and α-SMA (1:300). Sections were subsequently incubated for 2 h at room temperature with Alexa Fluor® 546-conjugated goat anti-mouse IgG and Alexa Fluor® 488-conjugated goat anti-rabbit IgG, each at a 1:500 dilution. Sections were mounted using VECTASHIELD® mounting medium containing DAPI and imaged using a Nikon fluorescence microscope.

For quantitative IHC/immunofluorescence analysis, five representative fields per group were evaluated by manual counting of marker-positive epithelial cells.

### 2.5 Mammary epithelial cell isolation and flow cytometry

Fourth mammary glands were harvested from female offspring at PND40. Tissues were mechanically minced using a tissue chopper and enzymatically digested with collagenase and hyaluronidase for 2 h at 37 °C, as previously described [28]. The resulting organoid fraction was further dissociated by sequential digestion with trypsin and dispase/DNase I and filtered through a 40-µm cell strainer to generate single-cell suspensions. Isolated mammary epithelial cells (MECs) were used for flow cytometric and functional analyses. MECs were independently isolated from five mice per group, with each biological sample assayed in triplicate.

For flow cytometry, 1 × 10⁶ cells were subjected to Fc-receptor blocking with anti-CD16/CD32 and subsequently stained with antibodies against lineage and mammary epithelial cell-surface markers according to established protocols [28]. PE-conjugated antibodies against CD31, TER-119, and CD45 were used to exclude endothelial, erythroid, and hematopoietic lineage-positive cells, respectively. CD24 and CD49f expression was assessed using biotinylated anti-CD24 followed by streptavidin-APC and FITC-conjugated anti-CD49f. Dead cells were excluded using 7-aminoactinomycin D (7-AAD).

Flow cytometric gating and quantification were performed using FlowJo software. Mammary epithelial subpopulations were identified based on CD24 and CD49f expression as indicated in the corresponding figures. MEC preparations from five mice per experimental group were analyzed.

### 2.6 Colony-forming, mammosphere, and three-dimensional culture assays

For colony-forming cell (CFC) assays, single-cell suspensions of primary MECs were plated at a density of 4 × 10³ cells per 60-mm culture dish and maintained at 37 °C in a 5% CO₂ atmosphere. After 10 days, colonies were fixed and stained with Wright–Giemsa stain. Colonies were imaged using a Nikon SMZ745T stereomicroscope and Nikon Elements Imaging System software and quantified using ImageJ.

For mammosphere assays, isolated MECs were seeded at a density of 2.5 × 10⁴ cells per well in ultra-low-attachment 24-well plates. Cells were cultured in EpiCult-B Mouse Medium (STEMCELL Technologies) supplemented with 10 µg/mL insulin (Sigma), 1 µg/mL hydrocortisone (Sigma), 1× B-27 (Thermo Scientific), 20 ng/mL epidermal growth factor (EGF; STEMCELL Technologies), 20 ng/mL basic fibroblast growth factor (bFGF; STEMCELL Technologies), 4 µg/mL heparin (STEMCELL Technologies), and 50 µg/mL gentamicin. After 7 days, primary mammospheres >30 µm in diameter were counted and imaged. For secondary mammosphere formation, primary mammospheres were dissociated into single cells by trypsinization, and 1,000 cells were replated under identical culture conditions. Secondary mammospheres were counted and imaged after an additional 7 days.

For three-dimensional (3D) culture assays, MECs were seeded in Matrigel at a density of 1.5 × 10⁴ cells per well in 48-well plates and cultured for 10 days. Colonies were subsequently stained with crystal violet, imaged, and quantified using ImageJ.

MECs were independently isolated from five mice per group, with each biological sample assayed in triplicate.

### 2.7 Western blot analysis

Western blot analyses were performed using mammary tissue from three independent mice per group. Fourth and fifth mammary glands were collected at the indicated endpoints, snap-frozen in liquid nitrogen, and stored until protein extraction. Tissues were homogenized at 4 °C in lysis buffer, and total protein concentrations were determined using the bicinchoninic acid (BCA) assay. Equal amounts of protein (30 µg per sample) were separated by 10–12% SDS-PAGE and transferred to nitrocellulose membranes. Membranes were blocked for 1 h in TBS containing 0.1% Tween-20 and 5% nonfat dry milk and incubated with primary antibodies overnight at 4 °C. Following washing, membranes were incubated for 1 h with the appropriate HRP-conjugated anti-mouse or anti-rabbit secondary antibodies (Thermo Scientific, Waltham, MA, USA). Protein signals were detected using Amersham enhanced chemiluminescence (ECL) reagents (GE Healthcare, Barrington, IL, USA) and captured using a FluorChem E imaging system (ProteinSimple, San Jose, CA, USA). Band intensities were quantified using ImageJ.

### 2.8 Statistical analysis

Statistical analyses and graph generation were performed using GraphPad Prism 8. ImageJ was used for quantification of Western blot signals and colonies in CFC and 3D culture assays. Data are presented as mean ± SD. Comparisons between two experimental groups were performed using unpaired, two-tailed Student’s *t*-tests. A *p* value <0.05 was considered statistically significant. In figures, * indicates *p* < 0.05 and ** indicates *p* < 0.01.

## 3. Results

### 3.1 In utero alcohol exposure induces alterations in mammary gland morphogenesis and epithelial proliferation in C57BL/6 mice

To determine whether prenatal alcohol exposure alters mammary development in the absence of an oncogenic driver, mammary gland morphogenesis was examined in female C57BL/6 offspring following in utero alcohol exposure (IUE) from GD11 to GD19. Mammary glands were evaluated at PND40, during active pubertal development, and at PND70, when the gland had reached young adulthood.

At PND40, whole-mount analysis revealed more extensive ductal development in IUE-exposed mammary glands compared with controls, accompanied by a significant increase in the number of terminal end buds (TEBs) (Fig. 1A, B). Because TEBs are highly proliferative structures that drive pubertal ductal outgrowth, we next examined epithelial proliferation by Ki67 immunofluorescence. IUE-exposed glands showed a marked increase in Ki67-positive mammary epithelial cells within SMA-defined ductal structures (Fig. 1C, D), consistent with enhanced proliferative activity during pubertal morphogenesis.

**Figure 1.**
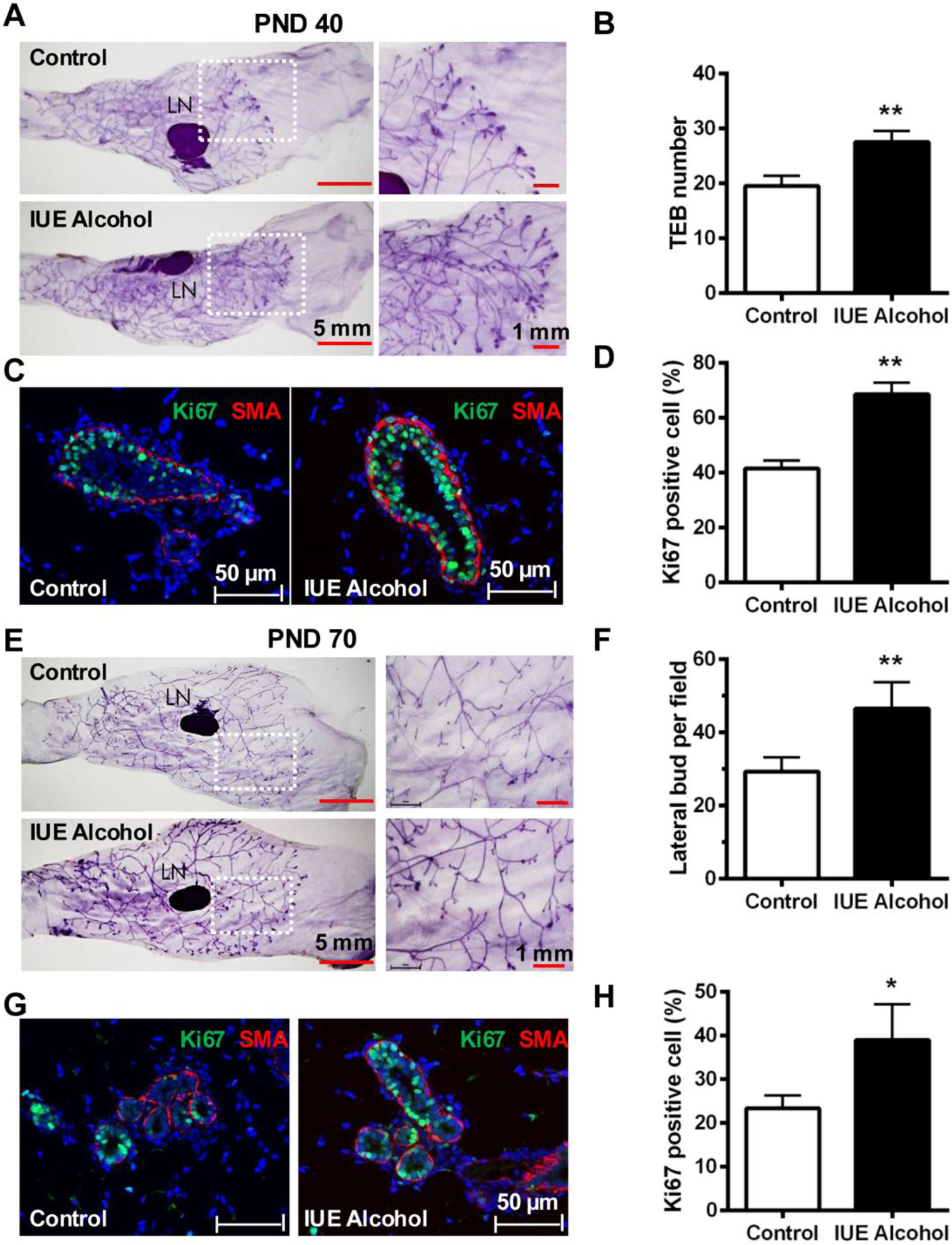
In utero alcohol exposure induces persistent alterations in mammary gland morphogenesis and epithelial proliferation. Female offspring of C57BL/6 dams exposed to control or 1.7% (v/v) alcohol liquid diet from gestational day (GD) 11 to GD 19 were examined at postnatal day (PND) 40 and PND70. **(A)** Representative whole-mount images of mammary glands at PND40. **(B)** Quantification of terminal end buds (TEBs) at PND40. **(C)** Representative immunofluorescence images of Ki67 (green) and α-SMA (red) in PND40 mammary gland sections. Nuclei were counterstained with DAPI (blue). **(D)** Quantification of Ki67-positive mammary epithelial cells at PND40. **(E)** Representative whole-mount images of mammary glands at PND70. **(F)** Quantification of lateral buds/branches per 10 mm² of mammary tissue at PND70. **(G)** Representative immunofluorescence images of Ki67 (green) and α-SMA (red) in PND70 mammary gland sections. Nuclei were counterstained with DAPI (blue). **(H)** Quantification of Ki67-positive mammary epithelial cells at PND70. Five representative fields per group were analyzed for Ki67 quantification. n = 5 mice per group. Data are presented as mean ± SD. \**p* < 0.05, \*\**p* < 0.01 versus control.

To determine whether these developmental alterations were transient or persisted beyond puberty, mammary glands were subsequently examined at PND70. Whole-mount analysis showed that IUE-exposed glands retained a more highly branched ductal architecture, with a significant increase in lateral buds/branches compared with age-matched controls (Fig. 1E, F). Increased epithelial proliferative activity also persisted at this stage, as demonstrated by a higher proportion of Ki67-positive mammary epithelial cells in IUE-exposed glands (Fig. 1G, H).

Together, these findings demonstrate that in utero alcohol exposure alters normal mammary gland developmental trajectories in a non-tumorigenic background, characterized by enhanced pubertal morphogenesis and epithelial proliferation that remain evident into young adulthood. The persistence of these phenotypes suggests that IUE produces more than a transient acceleration of pubertal development and prompted us to examine whether it is accompanied by sustained alterations in mammary epithelial cell composition and stem/progenitor activity.

### 3.2 In utero alcohol exposure alters mammary epithelial hierarchy and enhances stem/progenitor cell activity

Given the persistent morphogenetic and proliferative changes observed following IUE, we next asked whether prenatal alcohol exposure alters mammary epithelial cell composition and stem/progenitor activity. Mammary epithelial cells isolated from PND40 glands were analyzed by CD24/CD49f flow cytometry to assess major cell subpopulations and the MaSC-enriched mammary repopulating unit (MRU) fraction.

IUE significantly altered mammary epithelial composition, with expansion of the basal/myoepithelial compartment and a marked increase in the MRU-enriched population compared with controls, whereas the luminal compartment showed only a modest change (Fig. 2A–C). These findings indicate that prenatal alcohol exposure shifts the mammary epithelial hierarchy toward basal and stem/progenitor-enriched populations.

**Figure 2.**
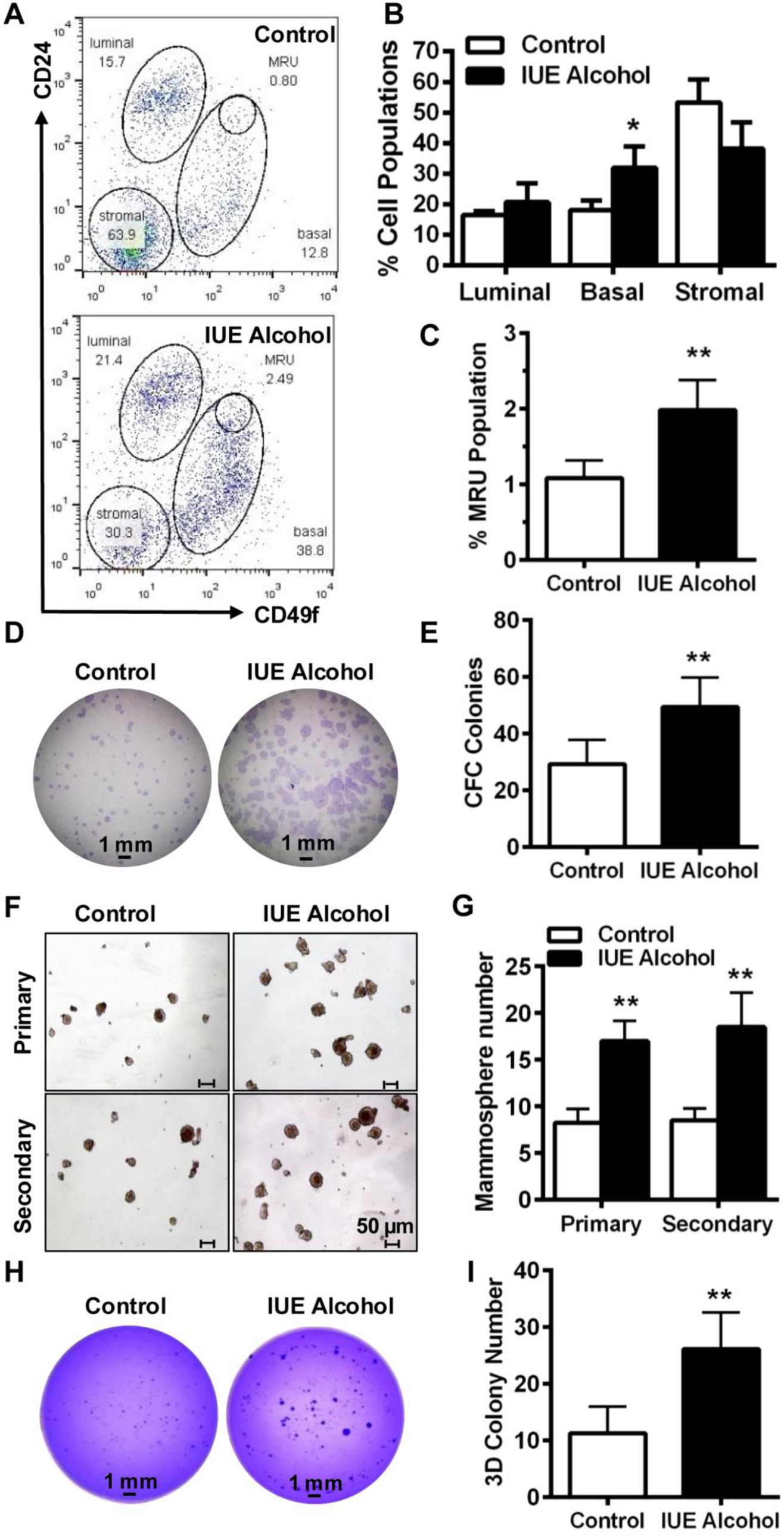
**In utero alcohol exposure alters mammary epithelial hierarchy and enhances stem/progenitor-associated activity. Primary mammary epithelial cells (MECs) were isolated from PND40 female C57BL/6 offspring exposed in utero to control or ethanol diet**. **(A)** Representative flow cytometry plots showing CD24/CD49f-defined mammary cell populations. The mammary repopulating unit (MRU)/MaSC-enriched population was identified as CD24^high^CD49f^high^. **(B)** Quantification of luminal, basal/myoepithelial, and stromal populations. **(C)** Quantification of the MRU/MaSC-enriched population. **(D)** Representative images of colony-forming cell (CFC) assays using primary MECs. **(E)** Quantification of colonies formed in CFC assays. **(F)** Representative images of primary and secondary mammospheres generated from control and IUE-exposed MECs. **(G)** Quantification of primary and secondary mammosphere formation. **(H)** Representative images of MEC colonies formed in 3D Matrigel culture after 10 days. **(I)** Quantification of colonies formed in 3D culture. n = 5 mice per group. Data are presented as mean ± SD. \**p* < 0.05, \*\**p* < 0.01 versus control.

To determine whether these population changes were accompanied by altered functional activity, primary MECs were evaluated using colony-formation cells (CFCs), mammosphere formation, and 3D culture assays. MECs from IUE-exposed mice generated significantly more colonies in CFC assays (Fig. 2D, E) and showed increased formation of both primary and secondary mammospheres (Fig. 2F, G), consistent with enhanced clonogenic and self-renewal-associated activity. In 3D Matrigel culture, IUE-derived MECs also formed significantly more colonies than control cells (Fig. 2H, I), indicating increased morphogenetic growth capacity.

Together, these findings demonstrate that IUE not only alters mammary epithelial composition but also enhances stem/progenitor-associated function. The concordance between MRU expansion and increased clonogenic, mammosphere, and 3D growth activities suggests persistent reprogramming of the mammary epithelial compartment, providing a cellular basis for the enhanced mammary development observed in vivo and prompting investigation of the signaling pathways associated with this phenotype.

### 3.3 In utero alcohol exposure enhances ERα signaling and associated proliferative and survival pathways

Given the enhanced epithelial proliferation and stem/progenitor activity following IUE, we next examined hormone-responsive signaling pathways involved in pubertal mammary development. At PND40, immunohistochemical analysis revealed increased p-ERα and Cyclin D1 expression in IUE-exposed mammary epithelium, with significant increases in both ducts and TEBs (Fig. 3A, B). The prominent changes within TEBs were consistent with the increased proliferative activity and TEB abundance observed following IUE.

**Figure 3.**
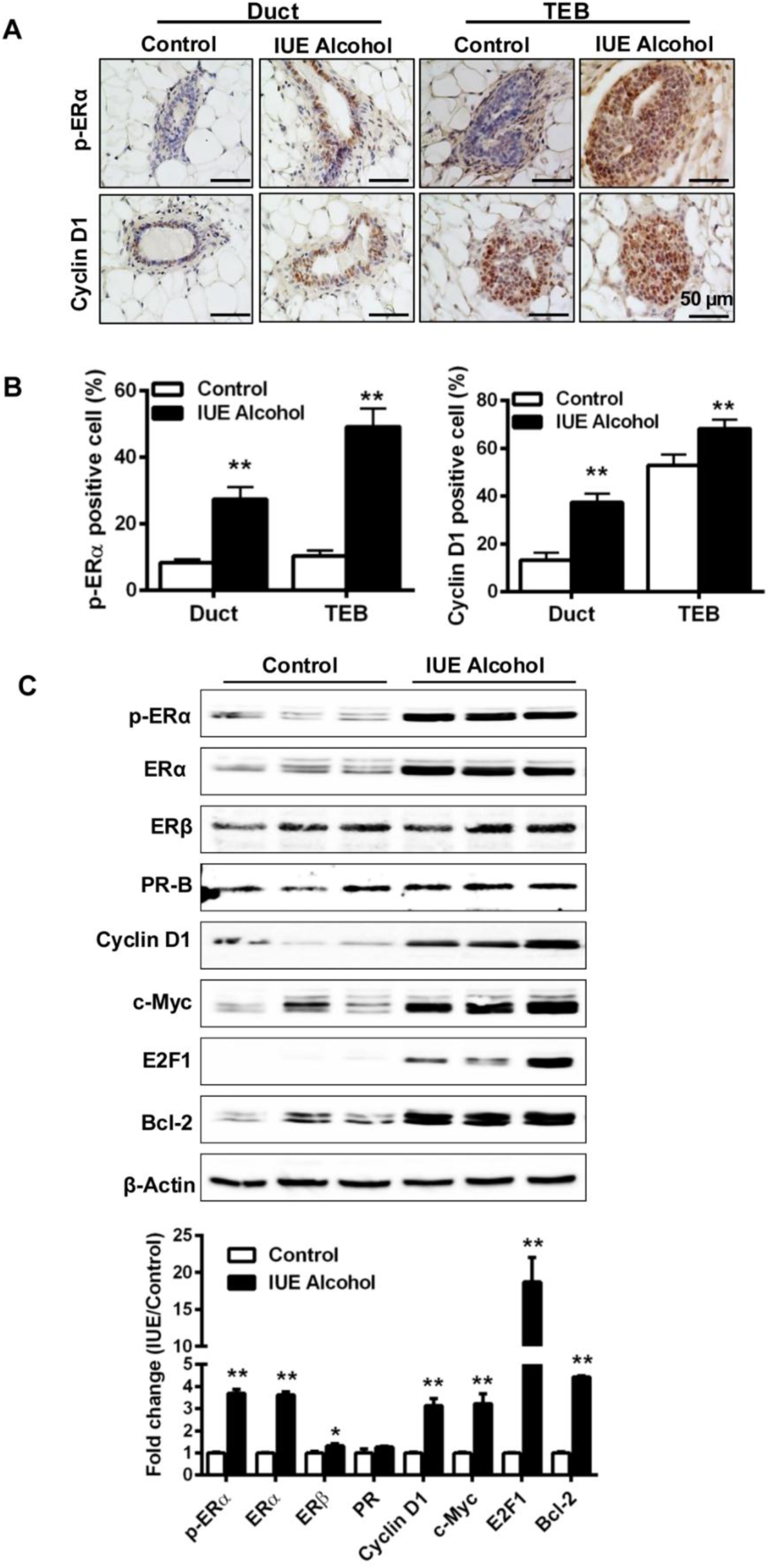
In utero alcohol exposure enhances ERα signaling and associated proliferative and survival pathways in pubertal mammary glands. Mammary glands were analyzed at PND40 following control or in utero alcohol exposure. **(A)** Representative immunohistochemical staining of phosphorylated ERα (p-ERα) and Cyclin D1 in mammary ducts and terminal end buds (TEBs). (B) Quantification of p-ERα- and Cyclin D1-positive epithelial cells in ductal and TEB regions. Five representative fields per group were analyzed. **(C)** Representative Western blots and densitometric quantification of ERα, p-ERα(Ser118), ERβ, PR-B, Cyclin D1, c-Myc, E2F1, and Bcl-2 in mammary gland lysates. β-Actin was used as a loading control. Western blot analyses were performed using mammary tissue from three mice per group. Data are presented as mean ± SD. \**p* < 0.05, \*\**p* < 0.01 versus control.

Western blot analysis further demonstrated increased ERα expression and phosphorylation at Ser118 in IUE-exposed mammary glands, whereas changes in ERβ and PR-B were comparatively modest (Fig. 3C). Enhanced ERα signaling was accompanied by increased expression of Cyclin D1, c-Myc, and E2F1, key regulators of cell-cycle progression, as well as the pro-survival protein Bcl-2 (Fig. 3C).

Together, these findings demonstrate that the developmental and cellular alterations induced by IUE are associated with sustained enhancement of ERα signaling and a coordinated proliferative and survival program in the pubertal mammary gland.

### 3.4 In utero alcohol exposure enhances growth, survival, and stemness-associated signaling

Because ER signaling interacts with multiple pathways controlling mammary epithelial growth and stem/progenitor activity, we next examined whether IUE was associated with broader changes in growth- and cytokine-responsive signaling at PND40. Western blot analysis demonstrated increased phosphorylation of AKT and ERK1/2 in IUE-exposed mammary glands, whereas total AKT and ERK2 levels remained relatively unchanged (Fig. 4A, B). These findings indicate enhanced activation of AKT and ERK signaling following prenatal alcohol exposure.

**Figure 4.**
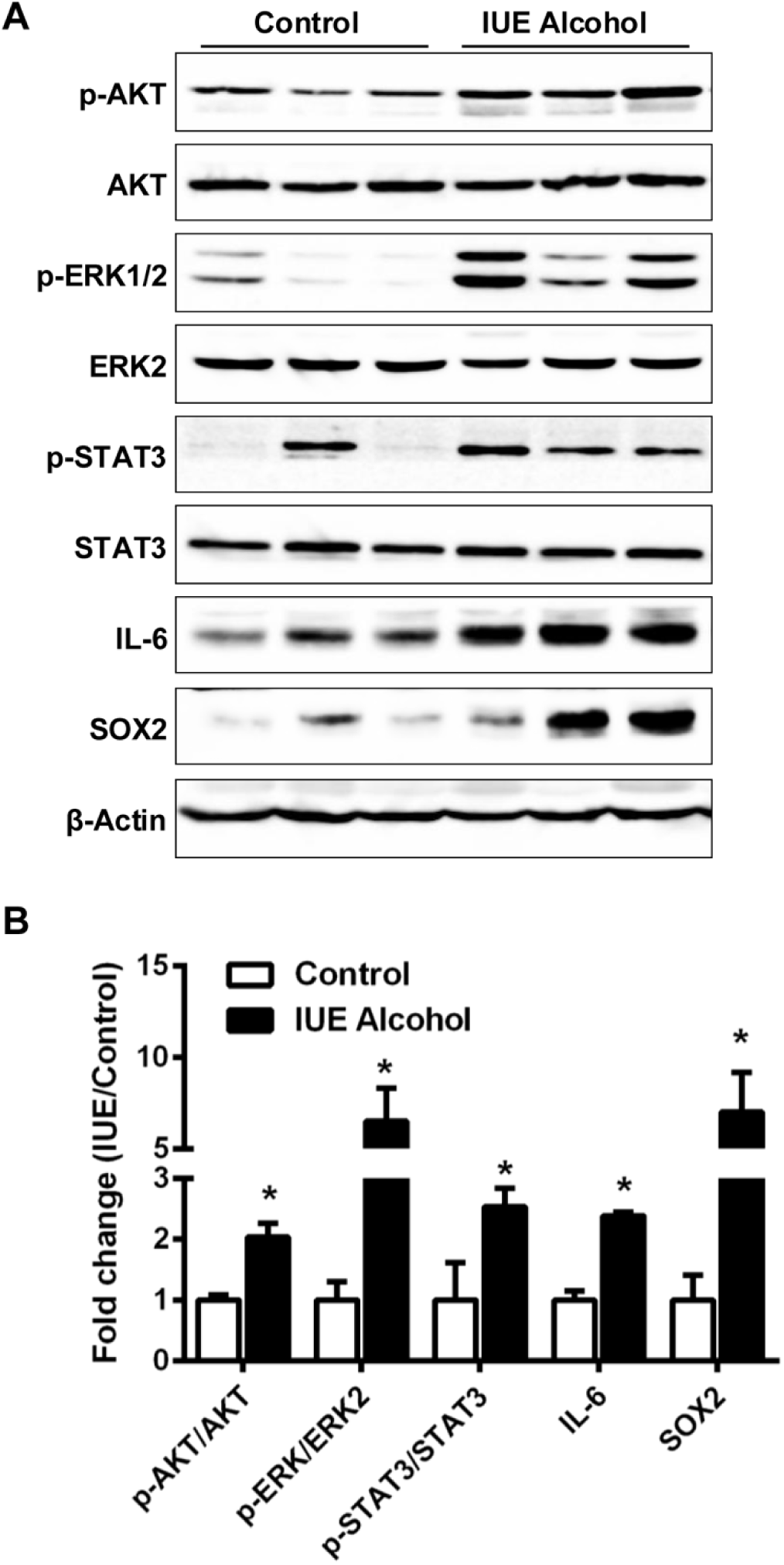
In utero alcohol exposure enhances growth, survival, and stemness-associated signaling in pubertal mammary glands. **(A)** Representative Western blots of p-AKT (Ser473) and total AKT, p-ERK1/2 (Thr202/Tyr204) and ERK2, p-STAT3 (Tyr705) and total STAT3, IL-6, and SOX2 in mammary gland lysates from control and IUE-exposed mice at PND40. β-Actin was used as a loading control. **(B)** Densitometric quantification of p-AKT/AKT, p-ERK1/2/ERK2, p-STAT3/STAT3, IL-6, and SOX2 expression relative to controls. Western blot analyses were performed using mammary tissue from three mice per group. Data are presented as mean ± SD. \**p* < 0.05, \*\**p* < 0.01 versus control.

IUE also increased STAT3 phosphorylation without a corresponding increase in total STAT3, accompanied by elevated IL-6 expression (Fig. 4A, B). In parallel, expression of the stemness-associated transcription factor SOX2 was markedly increased in IUE-exposed mammary glands. Thus, the enhanced stem/progenitor activity observed following IUE was associated with coordinated activation of AKT/ERK and IL-6/STAT3-associated signaling and increased SOX2 expression.

Together with the enhanced ERα signaling identified in Figure 3, these findings reveal a broader signaling state associated with the proliferative and stem/progenitor phenotypes induced by IUE. We therefore next asked whether these molecular alterations, like the developmental phenotype, persist beyond puberty into young adulthood.

### 3.5 In utero alcohol exposure induces persistent signaling alterations in adult mammary glands

Because the developmental phenotype induced by IUE persisted into young adulthood (Fig. 1), we next asked whether the associated molecular alterations were similarly sustained. At PND70, Western blot analysis demonstrated persistent increases in ERα expression and phosphorylation, accompanied by elevated Cyclin D1, E2F1, and Bcl-2 in IUE-exposed mammary glands (Fig. 5A). Increased p-ERα and Cyclin D1 expression was independently confirmed by immunohistochemistry, with significantly greater numbers of positive epithelial cells in IUE-exposed glands (Fig. 5B, C).

**Figure 5.**
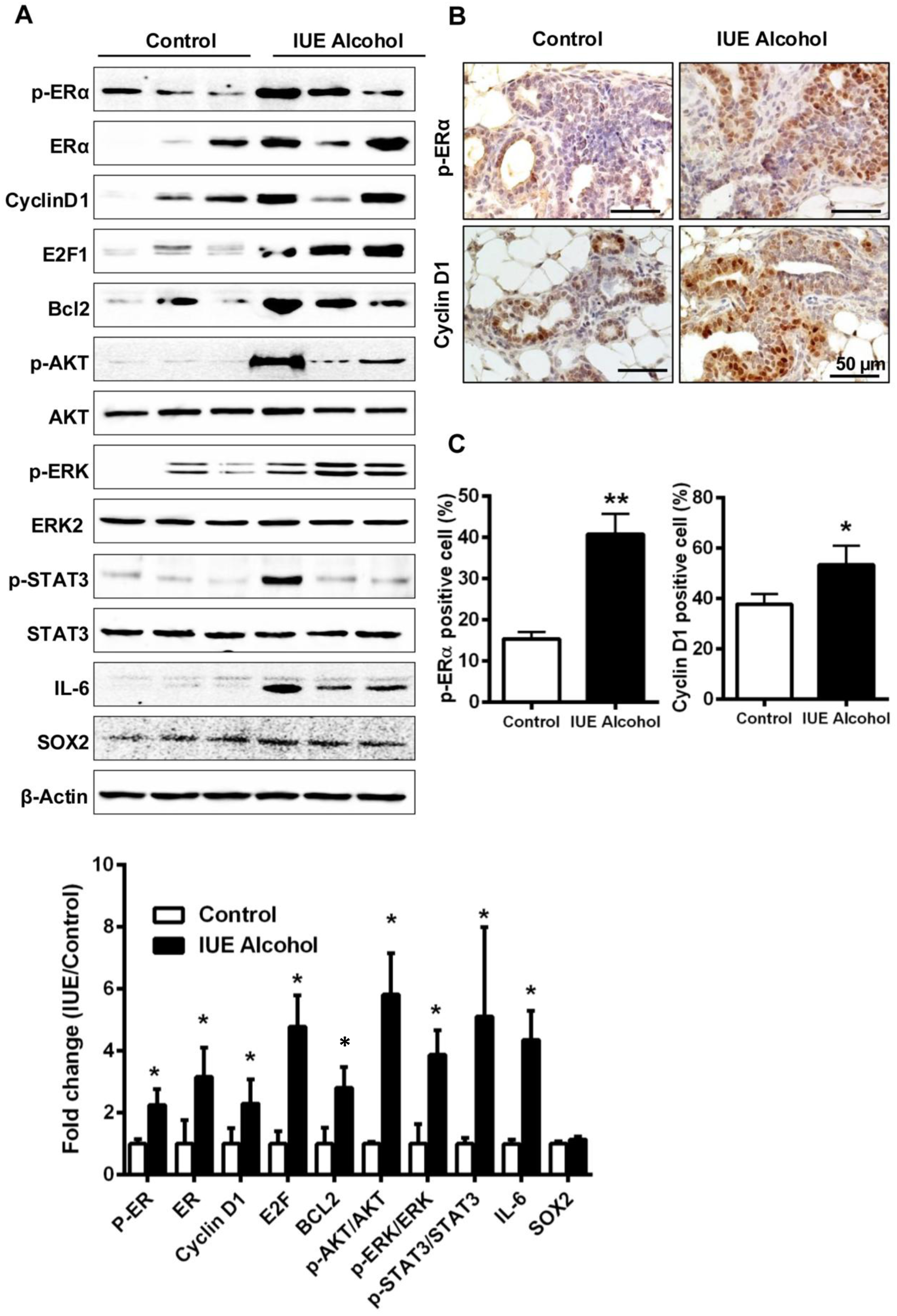
In utero alcohol exposure induces persistent signaling alterations in adult mammary glands. Mammary glands were analyzed at PND70 to determine whether signaling alterations associated with IUE persisted into young adulthood. **(A)** Representative Western blots and densitometric quantification of p-ERα, ERα, Cyclin D1, E2F1, Bcl-2, p-AKT (Ser473) and total AKT, p-ERK1/2 (Thr202/Tyr204) and ERK2, p-STAT3 (Tyr705) and total STAT3, IL-6, and SOX2. β-Actin was used as a loading control. Western blot analyses were performed using mammary tissue from three mice per group. **(B)** Representative immunohistochemical staining of p-ERα and Cyclin D1 in mammary gland sections from control and IUE-exposed mice at PND70. Scale bar, 50 µm. **(C)** Quantification of p-ERα and Cyclin D1-positive epithelial cells in mammary ducts. Five representative fields per group were analyzed. Data are presented as mean ± SD. \**p* < 0.05, \*\**p* < 0.01 versus control.

Activation of additional growth- and cytokine-responsive pathways also persisted into adulthood. IUE-exposed mammary glands exhibited increased phosphorylation of AKT, ERK, and STAT3, together with elevated IL-6 expression, whereas the corresponding total AKT, ERK2, and STAT3 levels remained relatively unchanged (Fig. 5A). In contrast to the pronounced increase observed at PND40, SOX2 expression was not substantially altered at PND70.

Together, these findings demonstrate that the effects of prenatal alcohol exposure extend beyond transient changes during pubertal mammary development. Persistent alterations in ERα-, AKT/ERK, and IL-6/STAT3-associated signaling parallel the sustained morphogenetic and proliferative phenotype observed at PND70, supporting long-term reprogramming of mammary gland development following IUE.

## 4. Discussion

The present study addresses an important question raised by previous studies of prenatal alcohol exposure and mammary tumor susceptibility: whether IUE produces any significant changes in mammary development in the absence of a subsequent carcinogenic or oncogenic challenge. Using non-tumorigenic C57BL/6 mice, we identified alterations in mammary morphogenesis, epithelial cell populations, and associated signaling pathways following prenatal alcohol exposure. The effects were evident during puberty as increased terminal end bud formation and epithelial proliferation and remained detectable in young adulthood as increased lateral branching and proliferative activity. These tissue-level changes were accompanied by shifts in mammary epithelial populations, enhanced stem/progenitor-associated activity, and coordinated changes in hormone-responsive, growth, and cytokine signaling. Thus, prenatal alcohol exposure appears to establish a persistent developmental state in the mammary gland rather than simply modifying the response of an already tumor-prone tissue.

This distinction is important in the context of earlier studies. Much of the experimental evidence linking prenatal alcohol exposure with mammary cancer risk has been obtained using carcinogen-induced or genetically tumor-prone models [20, 27]. These studies established that prenatal alcohol exposure can modify subsequent mammary tumor development, but the requirement for a carcinogenic challenge or oncogenic background made it difficult to determine which alterations preceded and which resulted from interactions with tumor-promoting signals. Our previous study using MMTV-erbB2 mice presented a similar question. IUE altered pubertal mammary morphogenesis, ER signaling, epithelial cell populations, and ErbB2-associated signaling and subsequently increased mammary tumor multiplicity [27]. The current results extend those observations by showing that several of these developmental and molecular phenotypes are also present in C57BL/6 mammary glands without ErbB2 overexpression. The altered mammary state produced by IUE therefore does not appear to require a pre-existing oncogenic driver, although such a state could influence how the tissue subsequently responds to oncogenic or environmental challenges.

Importantly, these developmental changes emerged during puberty and persisted into young adulthood. At PND40, IUE increased the number of TEBs, the highly proliferative epithelial structures responsible for pubertal ductal extension, together with increased Ki67-positive epithelial cells. By PND70, when pubertal ductal expansion has largely subsided, IUE-exposed glands continued to show increased lateral branching and epithelial proliferation. The phenotype therefore cannot be explained solely by an earlier onset or transient acceleration of pubertal development. Rather, prenatal exposure appears to alter the developmental trajectory of the gland in a manner that remains evident after the exposure itself, and the major period of pubertal ductal growth has passed. This persistent change in tissue organization is consistent with the broader concept of developmental programming, in which an early environmental exposure produces lasting changes in organ structure or regulatory responsiveness [29].

Changes in mammary epithelial populations provide a cellular correlation of this altered developmental trajectory [23]. At PND40, IUE increased the basal/myoepithelial compartment and the CD24^high^CD49f^high^ MRU population, which is enriched for mammary repopulating/stem cell activity. Importantly, the flow cytometric changes were accompanied by increased colony formation, primary and secondary mammosphere formation, and 3D growth of independently isolated mammary epithelial cells. The agreement among these assays argues that the population shifts are associated with altered functional properties rather than representing phenotypic marker changes alone. Because these assays measure overlapping but distinct aspects of progenitor activity, clonogenicity, and self-renewal-associated behavior, the data support an increase in stem/progenitor-associated activity following IUE. They do not, however, establish expansion of bona fide mammary stem cells in the transplantation-defined sense. Nevertheless, persistent alteration of these epithelial compartments provides a plausible cellular substrate through which a transient prenatal exposure could influence subsequent mammary development.

The molecular findings further suggest that this altered epithelial state is accompanied by changes in pathways that normally coordinate mammary growth and differentiation. ERα signaling was enhanced at PND40, with increased ERα expression and Ser118 phosphorylation together with increased Cyclin D1, c-Myc, E2F1, and Bcl-2. Increased p-ERα and Cyclin D1 within ducts and TEBs placed these changes in the epithelial structures undergoing active morphogenesis. ERα is a major regulator of pubertal mammary development [30], and its enhancement following IUE is therefore consistent with the increased epithelial proliferation and morphogenesis observed in vivo. Importantly, ER-associated alterations remained evident at PND70, when increased ERα/p-ERα, Cyclin D1, E2F1, and Bcl-2 accompanied the persistent morphological phenotype. These observations suggest that altered hormone responsiveness may be one component of the long-term developmental effect of prenatal alcohol exposure.

Signaling in EGFR family members and other receptor tyrosine kinases play a critical role in mammary development and tumor initiation [31]. IUE also affected signaling beyond the ER pathway. At PND40, increased phosphorylation of AKT and ERK occurred without comparable changes in total AKT or ERK, and increased STAT3 phosphorylation was accompanied by higher IL-6 expression. SOX2, a transcription factor associated with epithelial plasticity and stem/progenitor phenotypes [32], was also increased at this stage. These changes are consistent with the enhanced proliferative and stem/progenitor-associated activities observed in the same developmental period. Several components of this signaling pattern persisted at PND70, including increased AKT, ERK, and STAT3 phosphorylation and IL-6 expression. In contrast, the pronounced increase in SOX2 observed at PND40 was no longer apparent at PND70. This difference is informative because it suggests that IUE does not simply lock the mammary gland into a uniformly activated molecular state. Instead, some responses appear developmentally restricted, whereas others remain altered into adulthood.

The persistence of AKT and ERK activation in C57BL/6 mice also helps clarify observations from our earlier MMTV-erbB2 study. Because ErbB2 strongly engages these pathways, their activation in the transgenic model could have reflected an interaction between prenatal alcohol exposure and ErbB2 overexpression. Their elevation in the present non-tumorigenic model indicates that altered AKT/ERK signaling can occur independently of the ErbB2 transgene. Similarly, increased IL-6/STAT3 signaling provides an additional pathway associated with the IUE phenotype that is not dependent on an engineered oncogenic background. The present experiments do not identify the upstream signals responsible for AKT, ERK, or STAT3 activation, nor do they establish that these pathways cause the altered epithelial hierarchy. Rather, the combined data define a coordinated signaling context associated with the persistent developmental phenotype.

Taken together, the tissue, cellular, and molecular findings support a model in which prenatal alcohol exposure changes the developmental trajectory of the mammary gland at several interconnected levels. Enhanced pubertal morphogenesis occurs together with altered epithelial composition and increased stem/progenitor-associated activity, while ERα-, AKT/ERK-, and IL-6/STAT3-associated signaling provides a molecular context compatible with increased epithelial growth and plasticity. Importantly, major components of both the morphological and signaling phenotypes remain evident at PND70. In this framework, IUE is not required to directly initiate neoplastic transformation. Instead, developmental exposure may establish a mammary epithelial state with altered proliferative capacity, cellular composition, and responsiveness to regulatory signals, which could subsequently modify the tissue response to additional hormonal, environmental, or oncogenic influences.

This interpretation is consistent with the developmental origins of health and disease framework and may be particularly relevant to the mammary gland because its development extends well beyond the prenatal period. A developmental perturbation established during gestation can potentially influence subsequent rounds of hormonally driven epithelial expansion and remodeling during puberty and adult life. Previous carcinogen and tumor-prone studies suggest that prenatal alcohol exposure can modify later mammary tumor susceptibility. The present study does not directly test tumor development in C57BL/6 mice and therefore cannot establish that the developmental phenotype described here increases cancer risk. Rather, it identifies persistent changes in mammary architecture, epithelial stem/progenitor-associated activity, and signaling that are already present before the introduction of an experimental oncogenic stimulus. This distinction provides a basis for understanding how prenatal exposure and later tumor-promoting events might interact.

The present findings also raise questions about how these developmental changes are established and maintained after prenatal exposure. The association of ERα, AKT/ERK, and IL-6/STAT3 signaling with the altered epithelial phenotype provides several directions for determining which pathways contribute to these effects. Further studies at the cellular and lineage level, including molecular or epigenetic profiling, may also help define how prenatal exposure alters specific mammary epithelial populations and how these changes interact with the local tissue environment. Such studies will extend the current work from defining a persistent developmental phenotype toward understanding the processes that maintain it and influence subsequent mammary responses.

## 5. Conclusion

In summary, prenatal alcohol exposure altered mammary gland development in C57BL/6 mice in the absence of a carcinogenic challenge or genetically introduced oncogenic driver. IUE increased pubertal morphogenesis and epithelial proliferation, shifted mammary epithelial populations toward basal and MRU-enriched compartments, enhanced stem/progenitor-associated functions, and altered ERα-, AKT/ERK-, and IL-6/STAT3-associated signaling. Several of these morphological and molecular changes persisted into young adulthood. These results extend earlier tumor-model studies by showing that prenatal alcohol exposure produces persistent changes in normal mammary development in the absence of an oncogenic driver, providing a cellular and molecular context that may modify the response of the mammary gland to subsequent challenges.

## 6. Data Availability Statement

All data is contained within the manuscript.

## 7. Author Contributions

ZM: Data curation and analysis, writing-review & editing; AP: Data curation and analysis, writing-review & editing; EE: writing-review & editing; XY: Conceptualization, funding acquisition, investigation, project administration, writing – drafting, editing & review. All authors have read and agreed to the published version of the manuscript.

## 9. Acknowledgments

The authors extend their appreciation to the funding agency and support from colleagues in this department.

